# Generation of a white-albino phenotype from cobalt blue and yellow-albino trouts: inheritance pattern and chromatophores analysis

**DOI:** 10.1101/570861

**Authors:** Ricardo Shohei Hattori, Tulio Teruo Yoshinaga, Arno Juliano Butzge, Shoko Hattori-Ihara, Ricardo Yasuichi Tsukamoto, Neuza Sumico Takahashi, Yara Aiko Tabata

**Affiliations:** Salmonid Experimental Station at Campos do Jordão, UPD-CJ (APTA/SAA), Campos do Jordão; School of Veterinary Medicine and Animal Sciences, University of Sao Paulo, Department of Surgery, Sao Paulo; Department of Morphology, Institute of Bioscience of Botucatu, Sao Paulo State University, Botucatu; Department of Marine Biosciences, Tokyo University of Marine Science and Technology, Tokyo; Puriaqua Ltda., Sao Paulo; Sao Paulo Fisheries Institute (APTA/SAA), Sao Paulo

## Abstract

Albinism is the most common color variation described in fish and is characterized by a white or yellow phenotype according to the species. In rainbow trout *Oncorhynchus mykiss*, aside from yellow-albino phenotypes, cobalt blue variants with autosomal, recessive inheritance have also been reported. In this study, we investigated the inheritance pattern and chromatophores distribution/abundance of cobalt blue trouts obtained from a local fish farm. Based on crosses with wild-type and dominant yellow-albino lines, we could infer that cobalt blue are dominant over wild-type and co-dominant in relation to yellow-albino phenotype, resulting in a fourth phenotype: the white-albino. Analysis of chromatophores revealed that cobalt blue trouts present melanophores, as the wild-type, and a reduced number of xanthophores. As regards to the white-albino phenotype, they were not only devoid of melanophores but also presented a reduced number of xanthophores. Cobalt blue and white-albino trouts also presented a more elongated body shape and, most remarkably, a smaller pituitary gland compared to wild-type and yellow-albino, suggesting that the allele for blue color is somehow linked with this abnormal pituitary phenotype. These phenotypes represent interesting models for research on body pigmentation in salmonids and on the mechanisms behind endocrine control of color patterning.

## Introduction

The integumental pigmentary system of fish shares several conserved features with other vertebrate classes [1]. One of them is the presence of specific pigment-containing and light-reflecting cells involved mainly in body and eye pigmentation, known as chromatophores [2]. In the wild, body pigmentation is important for a variety of purposes such as protection against predators, advertising, sexual selection or warning [3]. In aquaculture industry, body pigmentation can be an economically relevant trait once external appearance can determine the commercial value of a given species, resembling the case of red Nile tilapia against wild-type [4] or the majority of freshwater/marine ornamental species [5]. In ecological studies, certain mutations in body coloration such as albinism can be very useful as visual markers for experimental tagging studies [6]. In genetic studies, they allow the evaluation of chromosome manipulation efficiency in gynogenic or triploid animals [7,8] and also for optimization of gene editing approaches [9,10].

In rainbow trout, different color variations have been described such as the albino, palomino, and cobalt/metallic blue phenotypes [11,12]. Regarding albinism, genes such as *tyr-1, tyr-2*, and *slc45a2* were shown to be implicated in pigmentation [9,10]. However, the mechanism or cells behind the blue phenotype is still unresolved. In majority of fish species, the blue color is caused by a light scattering phenomenon in purine and pteridine platelets in the iridophores. When combined with xanthophores other colors (e.g. green) can be generated whereas in their absence the blue color becomes prominent [5]. Indeed, a previous study reported a reduced number of xanthophores in blue trouts [13], suggesting that blue phenotype might be related to iridophores and the density of xanthophores in the skin.

The inheritance of albino phenotypes is determined mainly by recessive genes [11], but there is also a case of a dominant strain [14]. In case of blue phenotype, Blanc and collaborators [15] described a blue strain with autosomal, recessive inheritance. Although economically attractive, the recessive inheritance in such phenotypes may result in detrimental pleiotropic effects with regard to fertility and growth [15,16], once they require the presence of both alleles to be expressed. Such conditions, also promotes the expression of deleterious traits by means of recessive mutations. For instance, fries/juveniles of yellow-albino variants are reported as inferior in terms of survival and growth [11,16]. In case of cobalt blue, it was observed an association between this phenotype with obesity [17] and disorders in liver and kidney [18–20]. There are also reports showing complete sterility in both sexes due to the lack of a large part of the pituitary gland [19,21].

Under this scenario, this study aimed to better characterize our cobalt blue trouts by performing crosses with wild-type and dominant yellow-albino strains of rainbow trout. We first investigated the inheritance of body color in these and in another phenotype, named as white-albino. Then, we also compared the density of chromatophores, body growth, and pituitary morphology of these four phenotypes.

## Materials and Methods

### Ethics Statement

This study was conducted in accordance with animal rights of Colégio Brasileiro de Experimentação Animal (COBEA) and approved by Comitê de Ética em Experimentação Animal do Instituto de Pesca de São Paulo, Brazil (04/2017).

### Source of animals

A total of 137 rainbow trout juveniles with total body weight of about 100 g and cobalt blue phenotype were obtained from a local trout farm at Sapucaí-Mirim (Minas Gerais State, Brazil). According to the farmers, these cobalt blue rainbow trout juveniles were produced by crosses between cobalt blue and wild-type phenotypes. They were transferred to Salmonid Experimental Station at Campos do Jordão (SES-CJ) and reared for more 12 months until reaching sexual maturity for being used as broodstock in crosses experiments. Broodstock fish of wild-type and dominant yellow-albino strains from SES-CJ were used in crosses with cobalt blue females and males.

### Examination of color inheritance in crosses between different color phenotypes

To verify the inheritance of these phenotypes in relation to others, both females and males of cobalt blue (indicated as B+) rainbow trout were crossed between themselves and with the following strains: wild-type (bb) and dominant yellow-albino (AA). Gametes were manually stripped by applying gentle pressure along the abdomen and inseminated using a NaHCO_3_ solution (pH 8.0) for sperm activation. After two minutes, eggs were hydrated for 20 minutes and then washed with water. Egg incubation and larvae-juvenile rearing were performed separately in open flow through systems at temperatures ranging from 10 to 13 °C.

### Examination and measurement of chromatophores

Two-year-old fish with wild-type, yellow-albino, cobalt blue, and white-albino phenotypes were euthanized in an overdose of benzocaine solution After complete removal of scales, skin tissues with about 2 cm x 2 cm from the dorsal region located between the lateral line and dorsal fin. Skin samples were analyzed under a fluorescent microscope (BX53; Olympus, Japan). Images were captured with a CCD Camera (DP73) and analyzed by CellSens software (ver. 1.12; Olympus, Japan). The number of xanthophores from five to seven fields were counted for each color phenotype. The area of those cells (n = 15 cells per color phenotype) was measured using Image J software.

### Analysis of growth performances in four color phenotypes

Body weight (BW) and standard length (SL) were measured in four color phenotypes (n=14 to 25) obtained from the cross between blue female (F0) and white male (F1) All measurements were performed in 18 months-old adult females. Males were not included in the analysis due to the presence of precocious males in the four phenotypes. Condition factor (K) was calculated as BW (Kg)/SL (cm)^3^.

### Analysis of pituitary gland among four cobalt blue and wild-type phenotype

The pituitary gland was carefully dissected from juveniles (about 6 months) and sexually mature females and males (about 2 years) from the four color phenotypes (Table S1). The relative weight was calculated as pituitary weight (g) divided by total body weight (Kg).

### Statistical analyses

The significance of color phenotype deviations of the progenies from 1:1 and 3:1 was analyzed by the Chi-square test, with Microsoft Excel 2013 (Microsoft; Redmond, WA). The difference between groups was analyzed by One-way ANOVA followed Bonferroni’s test using GraphPad Prism (v.5.00; GraphPad Software, San Diego, CA, USA) or by Student’s T test. Differences were considered as significant for *p*<0.05.

## Results

### Color inheritance in crosses among blue, wild-type, and albino trouts

Crosses between blue female vs. wild-type male and wild-type female vs. blue male generated balanced proportions of both cobalt blue and wild-type phenotypes (Table 1). For three crosses between blue individuals (crosses #4, #5, and #6), the proportion between blue and wild-type was approximately 3:1, as supported by Chi-square test (Table 1). In the cross between blue female and dominant yellow-albino male, about half of the fish were yellow-albino as the mother while another half was white-albino (Figure 1A).

**Table 1.**
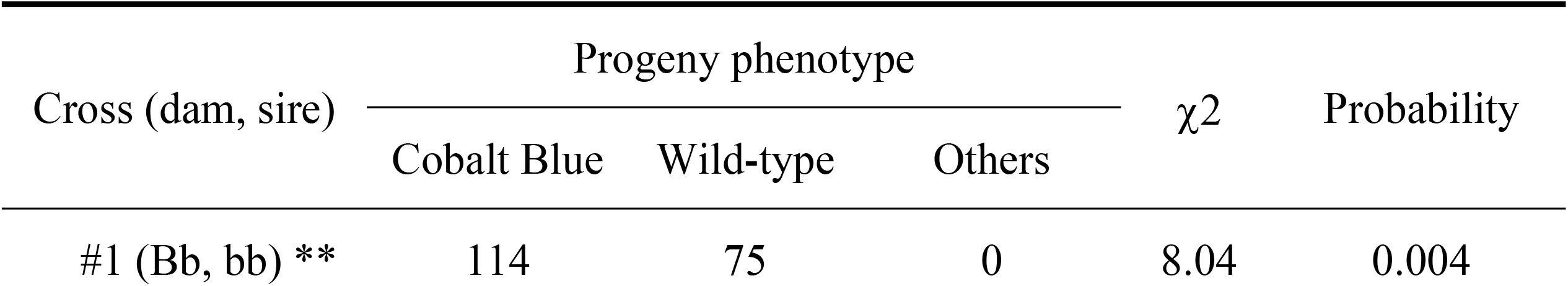
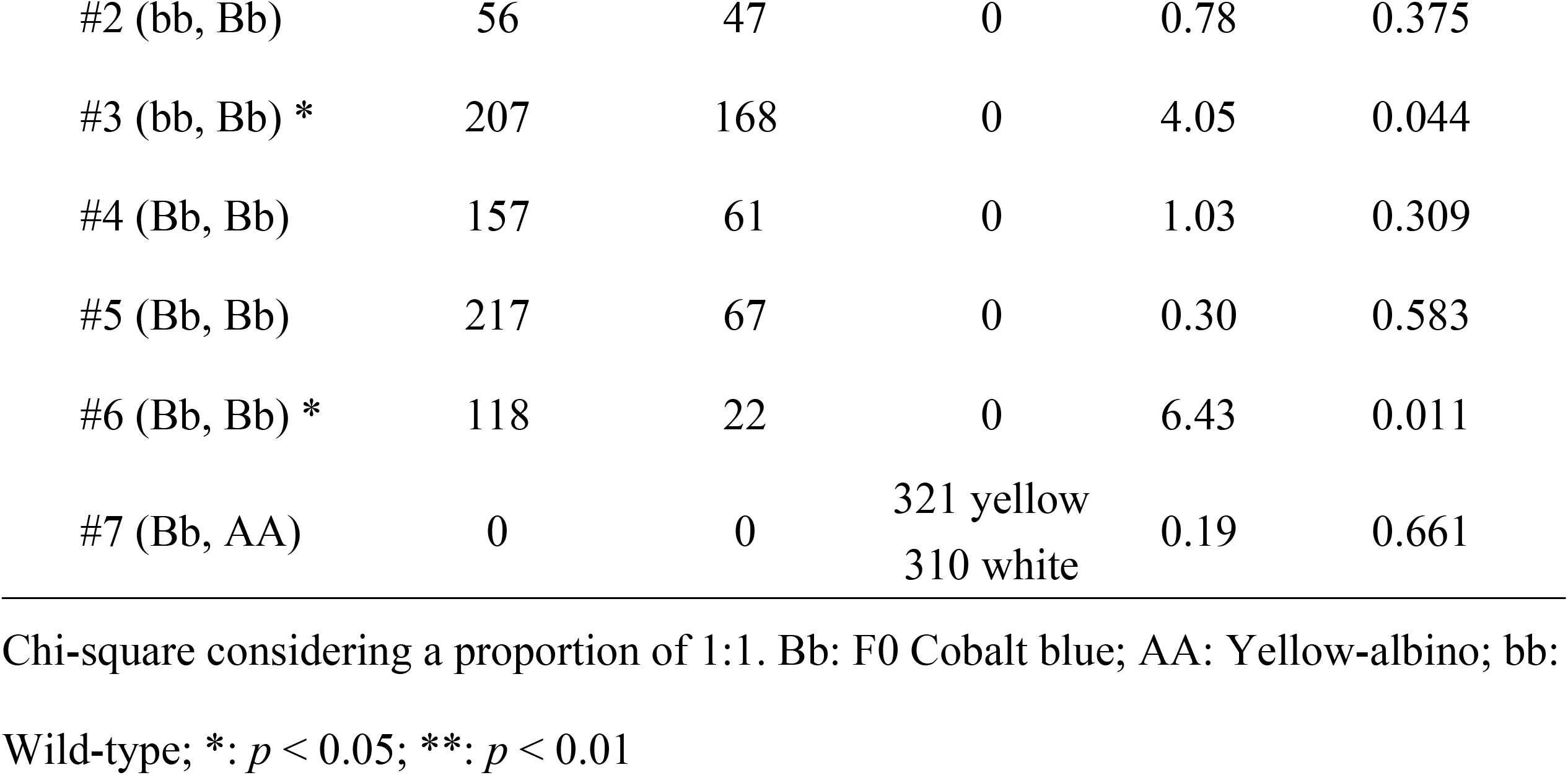
Frequency of color phenotypes generated by crosses using cobalt blue, wild-type, and yellow-albino (dominant) rainbow trout.

**Figure 1.**
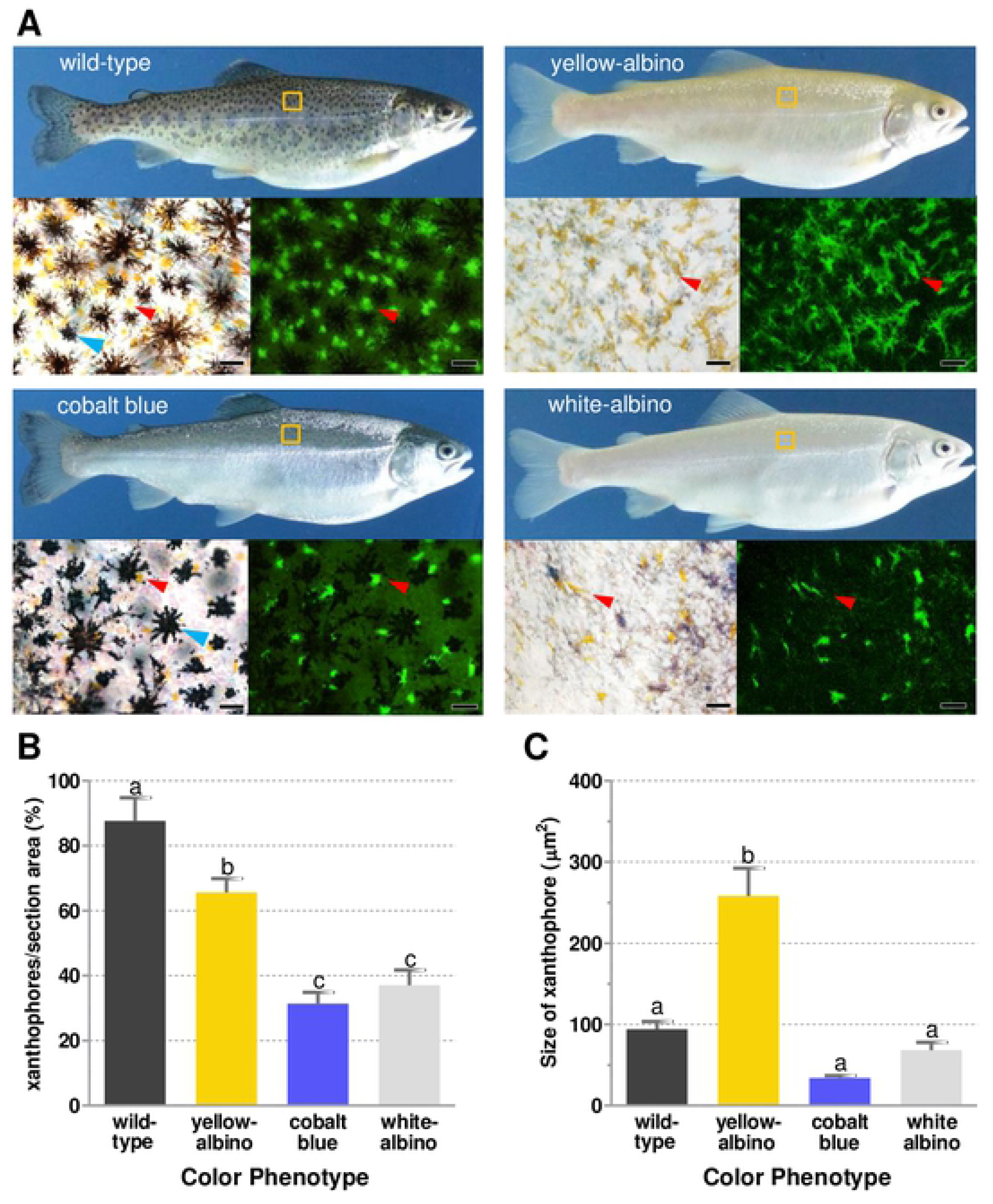
External appearance and chromatophores analysis in the four color phenotypes. **A**: Lateral view of wild-type, cobalt blue, yellow-albino, and white-albino female adults of rainbow trout, showing their respective chromatophores in bright field (left inferior) and under fluorescence (right inferior); the region selected for chromatophore analysis is depicted by a yellow box. Yellow- and white-albino are devoid of melanophores, which are indicated by blue arrow heads; yellow arrowhead indicates xanthophores. Scale bars indicate 25 μm. **B**: Average number and **C**: size of xanthophores in the dorsal region of trouts from each color phenotype (same as Figure 1A). Different letters represent statistically significant difference for *p*< 0.05 by one-way ANOVA analysis. Data are shown as average number ± SEM. Five to seven fields were randomly selected and analyzed for each color phenotype.

The F1 cobalt blue trout generated from crosses #4, #5, and #6 were used for progeny tests. Single-pair crosses between cobalt blue females (n=10) and wild-type males, and between wild-type females and cobalt blue males (n=10) generated eight all-cobalt blue progenies. The remaining 12 crosses yielded mixed-populations of blue and wild-type offspring (Table 2), whereby in six crosses the proportions did not deviated from 1:1 (*p* ≥ 0.05); in the remaining six, the proportion of wild-type were higher than that of cobalt blue in four crosses. From these results we can surmise that those eight individuals are homozygous for the blue color and the other 12 heterozygous for the *locus* responsible for the blue phenotype.

**Table 2.** Progeny tests using wild-type and F1 cobalt blue rainbow trouts derived from crosses between blue individuals (Crosses #4, #5, and #6 of Table 1).

| ID of parental<br>cobalt blue | Progeny phenotype | | $\chi^2$ | Probability |
| --- | --- | --- | --- | --- |
|  | Cobalt Blue | Wild-type |  |  |
| H1 | 159 | 149 | 0.32 | 0.568 |
| H2** | 137 | 243 | 29.56 | 0.000 |
| H3* | 186 | 149 | 4.08 | 0.043 |
| H4 | 296 | 0 | 296 | 0.000 |
| H5 | 161 | 175 | 0.58 | 0.455 |
| H6 | 290 | 0 | 290 | 0.000 |
| H7 | 229 | 0 | 229 | 0.000 |
| H8** | 93 | 172 | 23.55 | 0.000 |
| H9 | 29 | 0 | 29 | 0.000 |
| H10** | 70 | 106 | 7.36 | 0.006 |
| M1 | 398 | 0 | 398 | 0.000 |
| M2 | 18 | 0 | 18 | 0.000 |
| M3 | 159 | 188 | 2.42 | 0.119 |
| M4 | 108 | 111 | 0.04 | 0.839 |
| M5 | 42 | 39 | 0.11 | 0.738 |
| M6 | 45 | 48 | 0.09 | 0.755 |
| M7* | 67 | 95 | 4.83 | 0.027 |
| M8 | 67 | 0 | 67 | 0.000 |
| M9* | 48 | 26 | 6.54 | 0.010 |
| M10 | 104 | 0 | 104 | 0.000 |

Chi-square considering a proportion of 1:1. H: female; M: male; \*: $p < 0.05$ ; \*\*: $p < 0.01$

In a cross between F0 cobalt blue and F1 white-albino trouts, white-albino, cobalt blue, yellow-albino, and wild-type trouts were obtained (Table 3; Figure 2A) in a proportion that did not differ significantly from 3:3:1:1 (Chi-square test; *p* < 0.05). In the intercross between F1 white-albino trouts, white-albino, yellow-albino, cobalt blue, and wild-type trouts were obtained (Table 3; Figure 2B) in a proportion that did not differ significantly from 9:3:3:1 (Chi-square test; *p* < 0.05), respectively, as expected from a cross between double heterozygous, based on Mendelian segregation. The F1 yellow trouts were also intercrossed, yielding yellow and wild-type trouts (Table 3; Figure 2B) in a proportion that did not differ significantly from 3:1 (Chi-square test; *p* < 0.05), respectively, as expected from a cross between trouts with both alleles recessive for the blue *locus* (bb) and those for yellow in heterozygous condition (Aa), based on Mendelian segregation.

**Figure 2.**
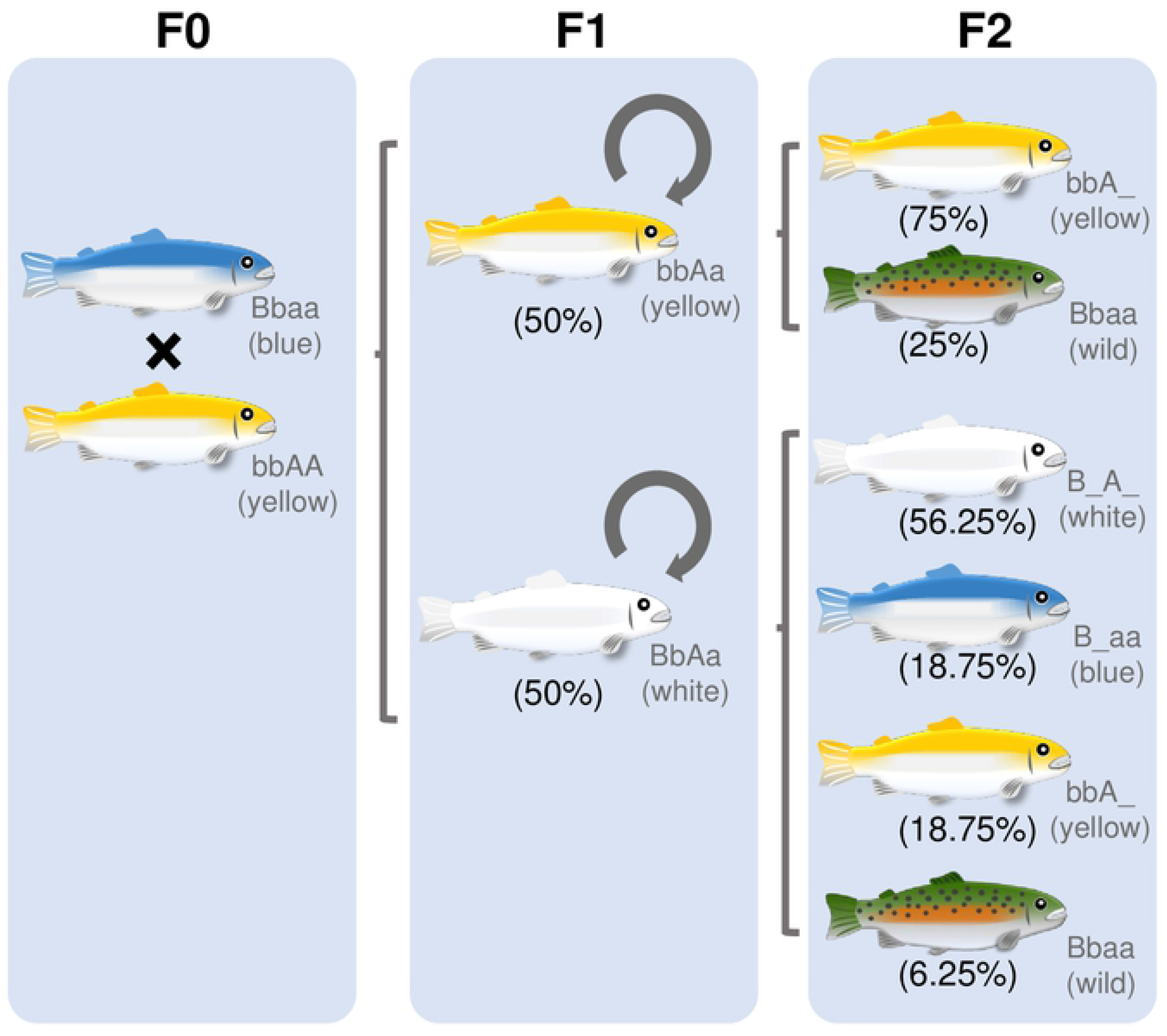
Schematic representation of crosses illustrating the pattern of color inheritance. A cobalt blue female and a yellow-albino male (F0) were initially crossed. F1 yellow- and white-albino trouts were crossed with the same color phenotype, as indicated by loops. The first group produced an F2 generation composed of yellow-albino and wild-type trouts in a proportion of 3:1, following Mendelian segregation for crosses between bbAa genotypes. Crosses between F1 white-albino produced an F2 generation composed of white-albino, cobalt blue, yellow-albino, and wild-type phenotypes in a 9:3:3:1 proportion, following Mendelian segregation for crosses between double heterozygous BbAa.

**Table 3.** Frequency of color phenotypes produced by a cross between F0 blue female and F1 male white-albino, and by intercrosses between F1 yellow-albino and F1 white-albino rainbow trout.

| Cross | dam<br>sire | Progeny phenotype | | | | $\chi^2$ | Probability |
| --- | --- | --- | --- | --- | --- | --- | --- |
|  |  | White-albino | Cobalt<br>Blue | Yellow<br>-albino | Wild-type |  |  |
| #8 | F0 Cobalt Blue<br>F1 White-albino | 157 | 165 | 75 | 64 | 7.760 | 0.051 |
| #9 | F1 Yellow-albino<br>F1 Yellow-albino | 0 | 0 | 185 | 60 | 0.034 | 0.853 |
| #10 | F1 White-albino<br>F1 White-albino | 379 | 126 | 132 | 44 | 1.075 | 0.783 |

Chi-square test considering proportions of 3:3:1:1 (#8), 0:0:3:1 (#9), and 9:3:3:1 (#10) of white, blue, yellow, and wild-type respectively; \*: $p < 0.05$ ; \*\*: $p < 0.01$

### Analysis of chromatophores in the four colors phenotypes

The analysis of melanophores demonstrated that these cells were present in wild-type and cobalt blue whereas in yellow- and white-albino rainbow trouts they were absent (Figure 1A; bright field images of chromatophores). In case of xanthophores (Figure 1A; fluorescent images), they were present in all phenotypes, including the white one. A comparison in the number of xanthophores (Figure 1B) revealed that they were most abundant in the wild-type, followed by dominant yellow-albino, cobalt blue, and white-albino phenotypes; between the last two there was no significant difference. The measurement of relative area of xanthophores in dominant yellow-albino trout were almost three times larger compared to other three color phenotypes (Figure 1C), that did not differ among themselves.

### Analysis of growth in four color phenotypes

The standard length was higher in the wild-type followed by the yellow albino, cobalt blue, and white-albino phenotypes (Figure 3A). A similar pattern was observed for body weight, whereby wild-type presented higher values in relation to yellow-albino, followed by cobalt blue and white-albino (Figure 3B). For standard length, white-albino showed significantly values lower than other phenotypes. For body weight, white-albino and cobalt blue showed significantly lower values (*p* < 0.05). In these phenotypes, we observed a more elongated body morphology in cobalt blue compared to the wild-type and yellow-albino trouts. This is supported by the condition factor (K) analysis and becomes more evident in animals with body length ranging from 30-35 cm (Figure 3C).

**Figure 3.**
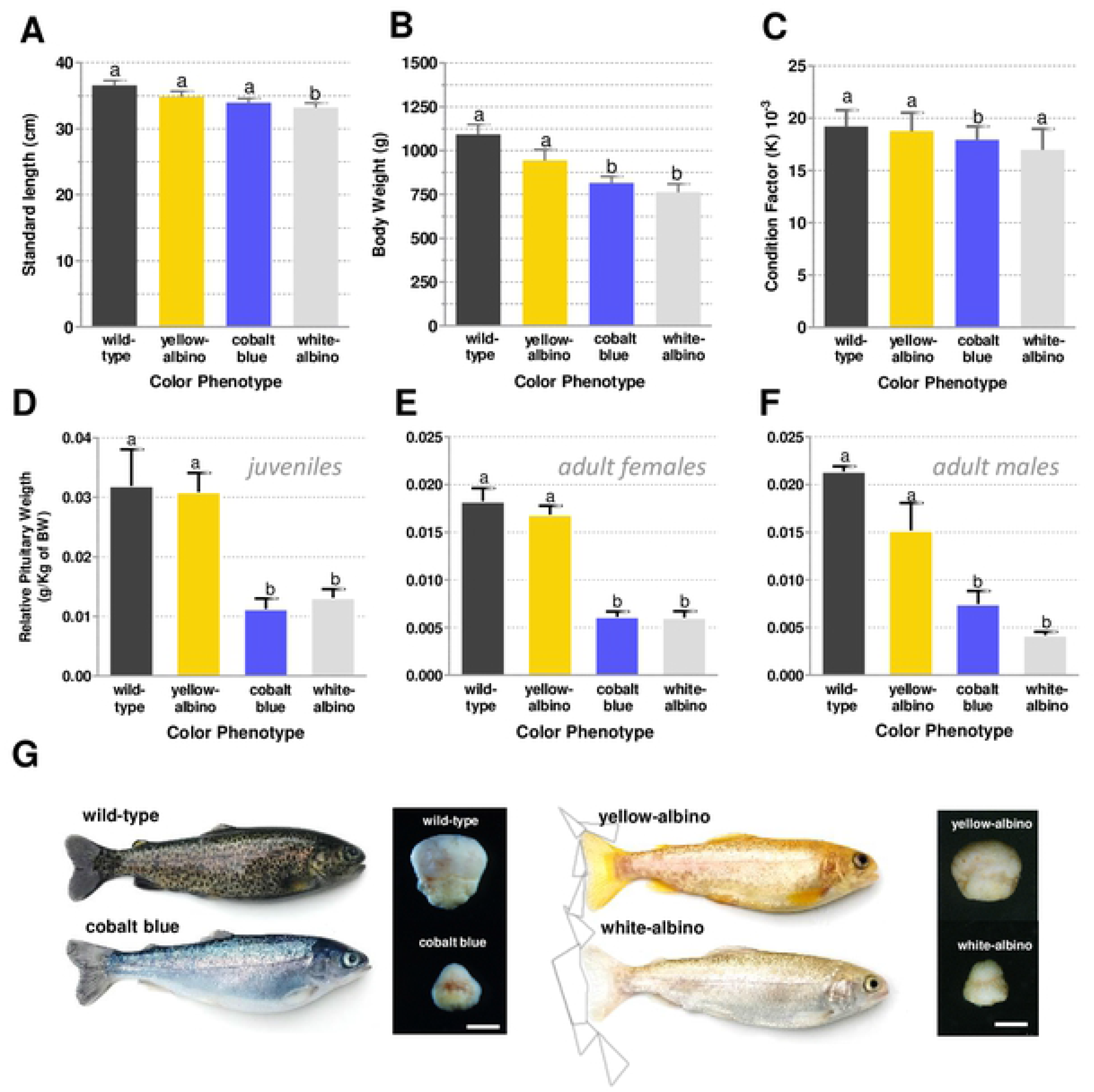
Comparative analysis of growth and pituitary morphology among color phenotypes. **A:** Standard length, **B:** body weight, and **C:** condition factor analysis in females from the four color phenotypes (n=14 *per* color phenotype). **D-F:** Relative pituitary weight in juveniles, adult females, and adult males from wild-type and cobalt blue trouts. **G:** Images of pituitary in wild-type, cobalt blue, yellow-albino, and white-albino juveniles with similar size and same age. Different letters and asterisk represent statistically significant difference by one-way ANOVA analysis for *p*< 0.05. Data are shown as average number ± SEM. Scale bar represents 0.5 mm.

### Analysis of pituitary in four color phenotypes

The comparison of pituitary morphology showed that juveniles and adults of cobalt blue and white-albino presented a smaller pituitary gland compared to wild-type and yellow-albino rainbow trout (Figure 3G). The values of relative pituitary weight of cobalt blue and white-albino phenotypes were significantly lower than those of wild-type and yellow-albino for both juveniles and adults (Figure 3D-F) (*p* < 0.05). No statistically significant differences were found between wild-type and yellow-albino as well as between cobalt blue and white-albino phenotypes.

## Discussion

Color variations from wild-type have been described for many fish species, especially in farmed ones submitted to intensive inbreeding [22]. In rainbow trout, color phenotypes including the yellow-albino and cobalt/metallic blue have been reported, but most of them display recessive inheritance and impairments related to growth or reproduction aspects [12,14]. In this study, we investigated the color inheritance and analyzed the density and types of chromatophores with focus in rainbow trout expressing a cobalt blue phenotype from juvenile to adult stages. Our results support that the allele responsible for the cobalt blue phenotype shows dominance over the wild-type and co-dominance in relation to yellow-albino; in the latter case a new, white-albino phenotype was generated. The analysis of chromatophores in cobalt blue revealed the presence of melanophores and a reduced number of xanthophores whereas the white-albino trouts were devoid of melanophores and, as the blue phenotype, presented lower density of xanthophores.

The blue phenotype represents an attractive phenotype due to external appearance with silvery colors [23], but the recessive inheritance was the main constraint for the establishment of commercial strains due to deficiencies in growth or reproduction [12,21,24]. The dominance of cobalt blue over the wild-type demonstrated by single-pair crosses represents an interesting characteristic, since the blue phenotype can be maintained even in a heterozygous condition, which thus allows the commercial production of hybrid blue strains by getting benefit from interstrain heterosis [25]. Our cobalt blue trout showed survival rates and fertility of both females and males comparable to the wild-type, but a slightly lower growth performances, which could be related to particular morphology of blue trouts (see discussion later). Nevertheless, the use of such a dominant strain, in association with wild-type strains displaying improved traits, may contribute to the development of new blue strains with high growth rates, tolerance to specific environmental conditions or resistance to pathogens [26–28].

The mechanisms involved in the blue coloration, especially in relation to chromatophores, in these trouts have not been clearly clarified. Although Blanc et al. [15] proposed that it is caused by the Tyndall effect, through a structural arrangement of melanin, the blue color in the majority of fish species is known to be caused by light scattering in platelets within iridophores [29,30]. The expression of blue color is enhanced when light is scattered by iridophore crystals (purines and pteridines) underlain by a dark surface, which could be the melanin of melanophores. When combined to yellow color from xanthophores, a green color is generated [5] whereas in its absence/reduction a blueish color is expressed [13]. The combined analysis of chromatophores and the results of crosses, especially between cobalt blue and yellow-albino trouts, are in agreement with this process, whereby a lower density of xanthophores was detected not only in the dorsal region, but also on the belly and undersides. Thus, besides the lower density of xanthophores, cobalt blue rainbow trout is supposed to require both iridophores and melanophores for the expression of blue color. On the other hand, the absence of melanophores and a lower abundance of xanthophores has probably yielded the white-albino phenotype, due to the reflectance of all colors by iridophores. The fact that blue and white-albino trouts do not acquire the characteristic reddish color in the lateral region and fins upon salmonization (Hattori, unpublished data), which is common in wild-type and yellow-albino, can be explained by low density of xanthophores, since carotenoids as astaxanthin are deposited in those chromatophores.

Melanophores, iridophores, and xanthophores are derived from neural crest cells, but the first two differentiate from a common pigment cell progenitor, whose fate is regulated by an interplay between *foxd3* and *mitfa* [31]. Xanthophores differentiate directly from neural crest cells or from an unknown precursor cell, presenting its own specification and differentiation pathway [32]. There are several genes involved in the differentiation and pigmentation of xanthophores described for fish, such as *pax7a*, *slc2a15b*, and *slc2a11b* [32], that could be implicated in the expression of blue color in our cobalt blue strain, but it seems that this color phenotype is caused by a mutation in a gene related to neuroendocrine pathway that somehow affect chromatophores rather than in a gene directly involved in chromatophore specification. This assumption is based on differences found in body morphology and also in the size of pituitary, that was inferior not only in cobalt blue but also in white-albino trouts that also carry the allele responsible for blue phenotype (B). Although a more detailed analysis is still required, particularly in terms of ontogeny, structure, and hormone secretion of pituitary, an intriguing aspect of cobalt blue trouts lies in their resemblance to the sea run form. This stage is characterized by a blueish color on dorsal region, silvery/whitish color on the belly and undersides, and an elongated body shape, which arises through the smoltification process during transition from freshwater to marine environment [33].

This process, characterized by physiological modifications, involves changes in body morphology and color patterning, which are triggered by environmental cues and mediated through a complex hormonal *milieu*, including the growth hormone/insulin-like growth factor 1, cortisol, thyroid hormones and also prolactin [34–36]. Based on the presence of many common traits between sea run form and the cobalt blue trouts, including a reduced number of xanthophores[37], it is possible that this phenotype could be caused by a mutation in a gene involved in the onset of smoltification process. Such mutation could lead to the appearance of a precocious “smolt-like” phenotype. A more detailed analysis on profiles of those hormones and in other physiological characteristics associated with smolts (e.g., Salt water-type chloride cells) could be performed to further explore the underlying mechanism behind the phenotype of blue trouts.

Compared to yellow-albino trouts, the white-albino is a rarer phenotype with few descriptions in fish species [32]. These two phenotypes can be obtained by mutations affecting melanophores or melanin synthesis [10,38,39]. Our crosses demonstrated that in rainbow trout, the white phenotype requires the presence of at least one dominant allele for each of blue and yellow-albino *loci* to be expressed. The analysis of chromatophores revealed that white-albino trouts were devoid of melanophores, as the yellow-albino phenotype, and presented a reduced number of xanthophores. In medaka, another population of chromatophores called leucophores responsible for the white color have been described, however these cells are absent in zebrafish and rainbow trout [32,40,41]. Thus, those information are suggestive that the white-albino phenotype could be simply explained by the concomitant absence of melanophores and xanthophores and the presence of iridophores that reflect all wavelengths resulting in a white-albino phenotype.

In conclusion, this study shows that the cobalt blue rainbow trout presents dominant inheritance in relation to wild-type and co-dominance with dominant yellow-albino trouts; in this last case a fourth phenotype called white-albino was generated. The allele responsible for the reduced number of xanthophores is supposedly a mutation in a gene related to neuroendocrine system based on differences observed in pituitary and body size and their involvement in chromatophore dynamics. Comparative studies among color variations in trout can be useful for endocrinology studies related to pigmentation process, particularly to ontogeny and specification of chromatophores, to growth performance, and also to pituitary development.

## Acknowledgments

This work was funded by Sao Paulo Research Foundation (FAPESP 2013/17612-9) to R.S.H. The authors thank L.R.S., A.D.S., and R.A.S.L. for fish rearing and maintenance.

## Author Contributions

Conceived and designed the experiments: RSH YAT. Performed the experiments: RSH YAT. Analyzed the data: RSH TTY AJB SHI YAT RYT. Wrote the paper: RSH TTY. Reviewed the manuscript for content: NST RYT.

---

**Table S1.** Standard length (cm; upper) and total body weight (Kg; bottom) of juveniles and adult fish used for pituitary comparison analysis. Data are represented by the average ± standard deviation. Numbers between brackets represent the sample size per color phenotype.

| Stage | Wild-type | Cobalt Blue | Yellow-albino | White-albino |
| --- | --- | --- | --- | --- |
| Juvenile (4) | 0.17±0.01 | 0.15±0.02 | 0.20±0.04 | 0.23±0.05 |
| | 20.3±0.7 | 20.3±0.7 | 20.0±1.2 | 20.5±1.6 |
| Adult female (3) | 0.98±0.06 | 1.02±0.03 | 1.24±0.12 | 1±0.16 |
|  | 36.5±1.5 | 36.5±1.9 | 36.2±1.2 | 34.7±1.9 |
| Adult male (3) | 0.33±0.07 | 0.4±0.15 | 1.07±0.08 | 0.89±0.42 |
|  | 24±1.3 | 25.0±2.6 | 35.5±0.5 | 31.1±5.5 |

